# The landscape of selection in 551 Esophageal Adenocarcinomas defines genomic biomarkers for the clinic

**DOI:** 10.1101/310029

**Authors:** AM Frankell, S Jammula, X Li, G Contino, S Killcoyne, S Abbas, J Perner, L Bower, G Devonshire, E Ococks, N Grehan, J Mok, M O’Donovan, S MacRae, M Eldridge, S Tavare, RC Fitzgerald, the Oesophageal Cancer Clinical and Molecular Stratification (OCCAMS) Consortium

**Affiliations:** MRC cancer unit, Hutchison/MRC research centre, University of Cambridge, Cambridge, UK; CRUK Cambridge institute, University of Cambridge, Cambridge, UK.; European Molecular Biology Laboratory, European Bioinformatics Institute (EMBL-EBI), Hinxton, UK; Department of Histopathology, Cambridge University Hospital NHS Trust, Cambridge, UK; Medical Research Council Cancer Unit, Hutchison/Medical Research Council Research Centre, University of Cambridge, Cambridge, UK; Cancer Research UK Cambridge Institute, University of Cambridge, Cambridge, UK; Department of Histopathology, Addenbrooke’s Hospital, Cambridge, UK; Oxford ComLab, University of Oxford, UK, OX1 2JD; Department of Computer Science, University of Oxford, UK, OX1 3QD; Cambridge University Hospitals NHS Foundation Trust, Cambridge, UK, CB2 0QQ; Salford Royal NHS Foundation Trust, Salford, UK, M6 8HD; Wigan and Leigh NHS Foundation Trust, Wigan, Manchester, UK, WN1 2NN; Royal Surrey County Hospital NHS Foundation Trust, Guildford, UK, GU2 7XX; Edinburgh Royal Infirmary, Edinburgh, UK, EH16 4SA; University Hospitals Birmingham NHS Foundation Trust, Birmingham, UK, B15 2GW; University Hospital Southampton NHS Foundation Trust, Southampton, UK, SO16 6YD; Cancer Sciences Division, University of Southampton, Southampton, UK, SO17 1BJ; Faculty of Medical and Human Sciences, University of Manchester, UK, M13 9PL; Gloucester Royal Hospital, Gloucester, UK, GL1 3NN; Guy’s and St Thomas’s NHS Foundation Trust, London, UK, SE1 7EH; Plymouth Hospitals NHS Trust, Plymouth, UK, PL6 8DH; Norfolk and Norwich University Hospital NHS Foundation Trust, Norwich, UK, NR4 7UY; Nottingham University Hospitals NHS Trust, Nottingham, UK, NG7 2UH; University College London, London, UK, WC1E 6BT; Norfolk and Waveney Cellular Pathology Network, Norwich, UK, NR4 7UY; Wythenshawe Hospital, Manchester, UK, M23 9LT; Edinburgh University, Edinburgh, UK, EH8 9YL; King’s College London, London, UK, WC2R 2LS; Karolinska Institutet, Stockholm, Sweden, SE-171 77; University Hospitals Coventry and Warwickshire NHS, Trust, Coventry, UK, CV2 2DX; Peterborough Hospitals NHS Trust, Peterborough City Hospital, Peterborough, UK, PE3 9GZ; Institute of Cancer and Genomic sciences, University of Birmingham, B15 2TT; GI science centre, University of Manchester, UK, M13 9PL.; Queen’s Medical Centre, University of Nottingham, Nottingham, UK, NG7 2UH; Department of Surgery and Cancer, Imperial College London, UK, W2 1NY; Queen’s Medical Centre, University of Nottingham, Nottingham, UK; Heart of England NHS Foundation Trust, Birmingham, UK, B9 5SS.; Centre for Cancer Research and Cell Biology, Queen’s University Belfast, Northern Ireland, UK, BT7 1NN.

## Abstract

Esophageal Adenocarcinoma (EAC) is a poor prognosis cancer type with rapidly rising incidence. Our understanding of genetic events which drive EAC development is limited and there are few molecular biomarkers for prognostication or therapeutics. We have accumulated a cohort of 551 genomically characterised EACs (73% WGS and 27% WES) with clinical annotation and matched RNA-seq. Using a variety of driver gene detection methods, we discover 77 EAC driver genes (73% novel) and 21 non-coding driver elements (95% novel), and describe mutation and CNV types with specific functional impact. We identify a mean of 4.4 driver events per case derived from both copy number events and mutations. We compare driver mutation rates to the exome-wide mutational excess calculated using Non-synonymous vs Synonymous mutation rates (dNdS). We observe mutual exclusivity or co-occurrence of events within and between a number of EAC pathways (GATA factors, Core Cell cycle genes, TP53 regulators and the SWI/SNF complex) suggestive of important functional relationships. These driver variants correlate with tumour differentiation, sex and prognosis. Poor prognostic indicators (SMAD4, GATA4) are verified in independent cohorts with significant predictive value. Over 50% of EACs contain sensitising events for CDK4/6 inhibitors which are highly correlated with clinically relevant sensitivity in a panel EAC cell lines and organoids.

## Introduction

Esophageal cancer is the eighth most common form of cancer world-wide and the sixth most common cause of cancer related death^1^. Esophageal Adenocarcinoma (EAC) is the predominant subtype in the west, including the UK and the US. The incidence of EAC in such countries has been rapidly rising, with a seven-fold increase in incidence over the last 35 years in the US^2^. EAC is a highly aggressive neoplasm, usually presenting at a late stage and is generally resistant to chemotherapy, leading to five-year survival rates below 15%^3^. It is characterised by very high mutation rates in comparison to other cancer types^4^ but also, paradoxically, there is a paucity of recurrently mutated genes. EACs also display dramatic chromosomal instability and thus may be classified as a C-type neoplasm which may be driven mainly by structural variation rather than mutations^5,6^. Currently our understanding of precisely which genetic events drive the development of EAC is highly limited and consequentially there is a paucity of molecular biomarkers for prognosis or targeted therapeutics available in the clinic.

Driver events undergoing positive selection during cancer evolution are a small proportion of total number of genetic events that occur in each tumour^7^. Methods to differentiate driver mutations from passenger mutations use features associated with known driver events to detect regions of the genome, often genes, in which mutations are enriched for these features^8^. The simplest of these features is the tendency of a mutation to co-occur with other mutations in the same gene at a high frequency, as detected by MutsigCV^9^. MutsigCV has been applied on several occasions to EAC cohorts^6,10,11^ and has identified ten known cancer genes as high confidence EAC drivers (TP53, CDKN2A, SMAD4, ARID1A, ERBB2, KRAS, PIK3CA, SMARCA4, CTNNB1 and FBXW7). Analysis of the non-coding genome has been performed by the PCAWG ICGC analysis and identified a significantly mutated enhancer associated with TP53TG1^12^. However these analyses leave most EAC cases with only one known driver mutation, usually TP53, due to the low frequency at which other drivers occur. Equivalent analyses in other cancer types have identified three or four drivers per case^13,14^. Similarly, detection of copy number driver events in EAC has relied on identifying regions of the genome recurrently deleted or amplified, as detected by GISTIC^10,15-18^. However, GISTIC identifies relatively large regions of the genome, often containing hundreds of genes, with little indication of which specific gene-copy number aberrations (CNAs) may actually confer a selective advantage. There are also several non-selection based mechanisms which can cause recurrent CNAs, such as fragile sites where a low density of DNA replication origins causes frequent structural events at a particular loci. These have not been differentiated properly from selection based recurrent CNAs^19^. Epigenetic events, for example methylation, may also be important sources of driver events in EAC but are much more difficult to assess formally for selection.

Without proper annotation of the genomic variants which drive the biology of EAC tumours we are left with a very large number of events, most of which are likely to be inconsequential, making it extremely difficult to detect statistical associations between genomic variants and various biological and clinical parameters. To address these issues, we have accumulated a cohort of 551 genomically characterised EACs using our esophageal ICGC project, which have high quality clinical annotation, associated whole genome sequencing (WGS) and RNA-seq on cases with sufficient material. We have augmented our ICGC WGS cohort with publically available whole exome^20^ and whole genome sequencing^21^ data. We have applied a number of complementary driver detection tools to this cohort, using a range of driver associated features combined with analyses of RNA expression to produce a comprehensive assessment and characterisation of mutations and CNAs under selection in EAC. We then use these events to define functional cell processes that have been selectively dysregulated in EAC and identify novel, clinically relevant biomarkers for prognostication, which we have verified in independent cohorts. Finally, we have used this compendium of EAC driver variants to provide an evidence base for targeted therapeutics, which we have tested *in vitro*.

## Results

### A Compendium of EAC driver events and their functional effects

In 551 EACs we called a total of 11,813,333 single nucleotide variants (SNVs) and small insertions or deletions (Indels), with a median of 6.4 such mutations / Mb (supplementary figure 1), and 286,965 copy number aberrations (CNAs). We also identified 134,697 structural variants (SVs) in WGS cases. Mutations or copy number variants under selection were detected using specific driver associated-mutation features (Fig 1A). We use several complementary driver detection tools to detect each feature, and each tool underwent quality control to ensure reliability of results (see methods). These features include highly recurrent mutations within a gene (dNdScv^22^, <u>ActivedriverWGS</u>^23^, MutsigCV2^9^), high functional impact mutations within a gene (OncodriveFM^24^, ActivedriverWGS^23^), mutation clustering (OncodriveClust^25^, eDriver^26^ and eDriver3D^27^) and recurrent amplification or deletion of genes (GISTIC^15^) undergoing concurrent over or under-expression (see methods) (Fig 1A)^8^.

**Figure 1.**
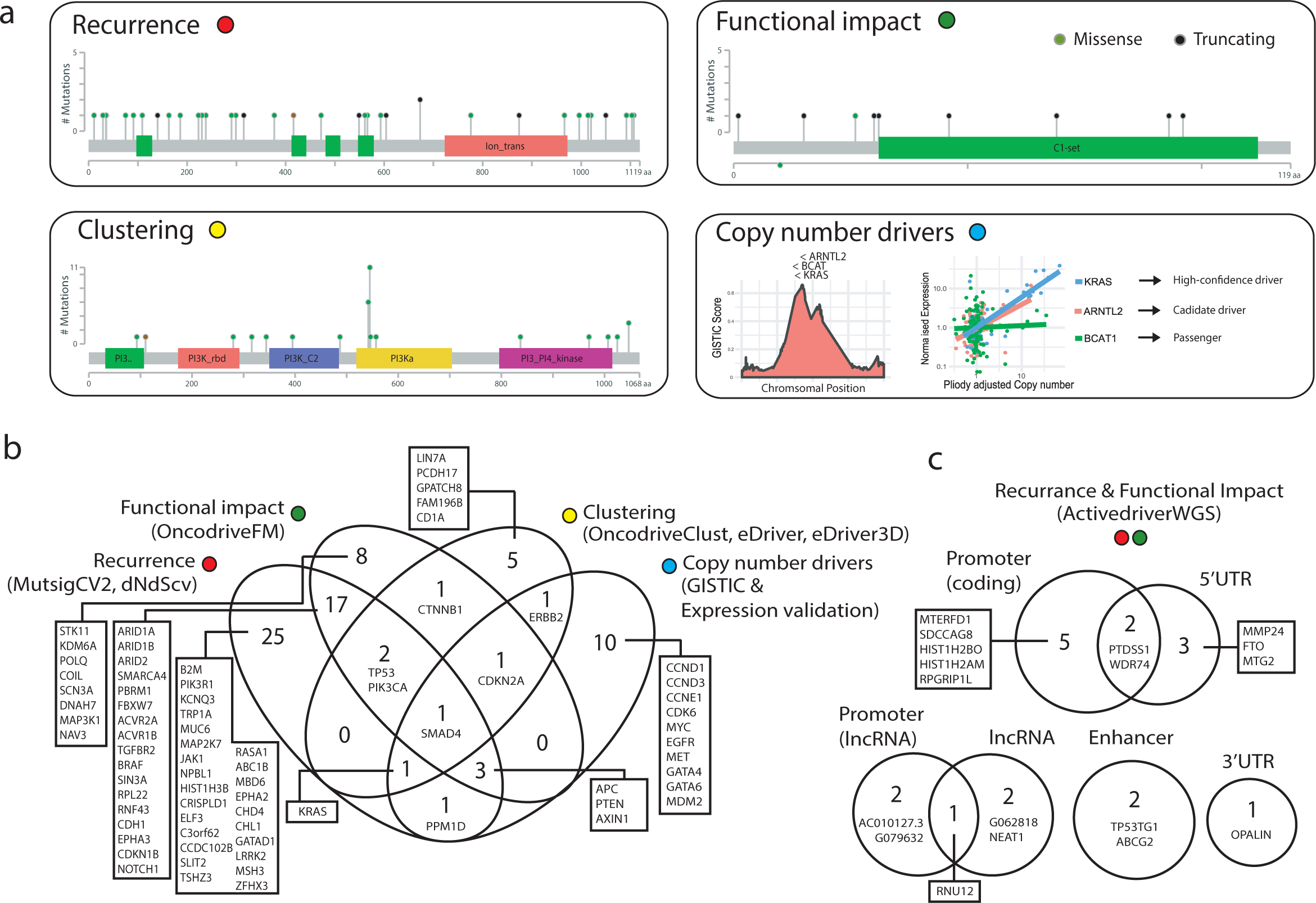
Detection of EAC driver Genes. **a.** Types of driver-associated features used to detect positive selection in mutations and copy number events with examples of genes containing such features **b.** Coding driver genes identified and their driver-associated features. **c.** Non-coding driver elements detected and their element types.

These complementary methods produced highly significant agreement in calling EAC driver genes, particularly within the same feature-type (supplementary figure 2) and on average more than half of the genes identified by one feature were also identified by other features (Fig 1B). In total seventy six EAC driver genes were discovered, 86% of which have not been detected in EAC previously^10,11,16-18,20^ and 69% are known drivers in pan-cancer analyses giving confidence in our methods^22,28,29^. To detect driver elements in the non-coding genome we used ActiveDriverWGS^23^ a recently benchmarked^30^ method using both function impact prediction and recurrence to determine driver status (Fig 1C, supplementary figure 3). We discovered 21 non-coding driver elements using this method. We have recovered several known non-coding driver elements from the pan-cancer PCAWG analysis^12^ including an enhancer on chr7 linked to TP53TG1, a gene required for TP53 action, the only non-coding driver found in EAC in PCAWG and the promoter/5’UTR regions of PTDSS1 and WRD74 which are novel in EAC but were found in other cancer types. We also identified completely novel non-coding cancer driver elements including in the 5’UTR of MMP24 and promoters of two related histones (HIST1H2BO and HIST1H2AM).

EAC is notable among cancer types for harbouring a high degree of chromosomal instability^21^. Using GISTIC we identified 149 recurrently deleted or amplified loci across the genome (Fig 2A). To determine which genes within these loci confer a selective advantage when they undergo CNAs we use a subset of 116 cases with matched RNA-seq to detect genes within these loci in which homozygous deletion or amplification causes a significant under or over-expression respectively, a prerequisite for selection of CNAs. The majority of genes in these regions showed no significant CN associated expression change (74%), although work in larger cohorts suggests we may be underpowered to detect small expression changes^31^. We observed highly significant expression changes in 17 known cancer genes within GISTIC peaks such as ERBB2, KRAS and SMAD4 which we designate high-confidence EAC drivers. We also found five tumour suppressor genes where copy number loss was not necessarily associated with expression modulation but tightly associated with presence of mutations leading to LOH, for example ARID1A and CDH11. CDH11 was not identified by our driver gene detection methods but this would suggest it may be a promising candidate for further validation. To determine whether copy number changes in genes not previously associated with cancer may contribute to oncogenesis we searched for genes with similar expression-CN profile as most of our high-confidence drivers (see methods). We found 140 such cases which we designated “candidate copy number (CN) drivers” (supplementary tables 1-4). Not all candidate drivers are likely to be true CN-drivers. However, several candidate drivers such as ZNF131, YES1 and PIBF1 are not accompanied by other drivers in their GISTIC peak and contain extrachromosomal-like events, hence are promising candidates for further study.

**Figure 2.**
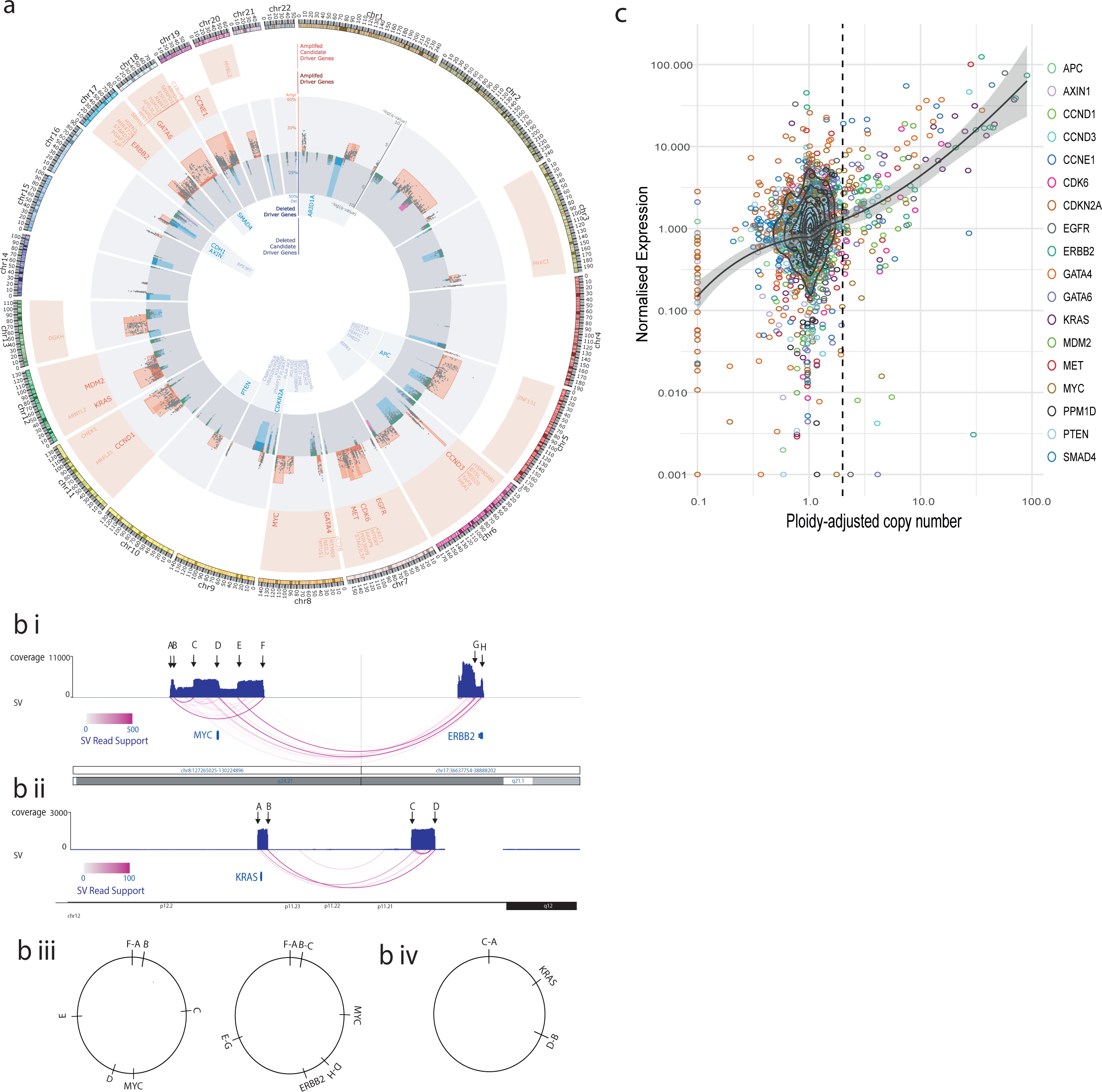
Copy number variation under positive selection. **a**. Recurrent copy number changes across the genome identified by GISTIC. Frequency of different CNV types are indicated as well as the position of CNV high confidence driver genes and candidate driver genes. The q value for expression correlation with amplification and homozygous deletion is shown for each gene within each amplification and deletion peaks respectively and occasions of significant association between LOH and mutation are indicated in green. Purple deletion peaks indicate fragile sites. **b**. Examples of Extrachromosomal-like amplifications suggested by very high read support SVs at the boundaries of highly amplified regions produced from a single copy number step. In the first example (bi) two populations of extrachromosomal DNA are apparent (biii), one amplifying only MYC and the second also incorporating ERBB2 from a different chromosome. In the second example (bii) an inversion has occurred before circularization and amplification around KRAS (biv). **c**. Relationship between copy number and expression in CN driver genes.

In a subset of GISTIC loci, we observed extremely high copy number amplification, commonly greater than 100 copies, and these loci were highly correlated with presence of CN-drivers (Ploidy adjusted Copy number >10, Wilcox test, p<Ex10^−6^) (supplementary figure 4). We use copy number adjusted ploidy to define amplifications as it produces superior correlation with expression data than absolute CN alone. Ploidy of our samples varies from 2-6 (3.5 on average) and hence Ploidy adjusted copy number of >10 cut off translates into >20-60 absolute copies (on average 35 copies). To discern a mechanism for these ultra-high amplifications we assessed structural variants (SVs) associated with these events and the copy number steps surrounding them. For many of these events the extreme amplification was produced largely from a single copy number step the edges of which were supported by structural variants with ultra-high read support. Two examples are shown in Fig 2B and further examples in supplementary figure 5. In the first example circularisation and amplification initially occurred around MYC but subsequently incorporated ERBB2 from an entirely different chromosome and in the second an inversion has been followed by circularisation and amplification of KRAS. A pattern of extrachromosomal amplification via double minutes has been previously noted in EAC^21^, and hence we refer to this amplification class with ultra-high amplification (Ploidy adjusted Copy number >10) as ‘extrachromosomal-like’. Several deletion loci co-align with fragile sites (Fig 2A). Most deletion loci were dominated by heterozygous deletions while a small subset had a far higher percentage of homozygous deletions including CDKN2A and several associated with fragile site loci (Fig 2A). For some cases we may have been unable to identify drivers in loci simply because the aberrations do not occur in the smaller RNA-seq matched cohort.

We found extrachromosomal-like amplifications had an extreme and highly penetrant effects on expression while moderate amplification (ploidy adjusted copy number > 2) and homozygous deletion had highly significant (Wilcox test, p<Ex10^−4^ and p<Ex10^−3^ respectively) but less dramatic effects on expression with a lower penetrance (Fig 2C). This lack of penetrance was associated with low cellularity (fisher’s exact test, expression cut off = 2.5 normalised FPKM, p<0.01) in amplified cases but also likely reflects that genetic mechanisms other than gene-dosage can modulate expression in a rearranged genome. We also detected several cases of over expression or complete expression loss without associated CN changes which may reflect non-genetic mechanisms for driver dysregulation. For example, one case overexpressed ERBB2 at 28-fold median expression however had entirely diploid CN in and surrounding ERBB2 and a second case contained almost complete loss of SMAD4 expression (0.008-fold median expression) despite possessing 5 copies of SMAD4.

### Landscape of driver Events in EAC

The overall landscape of driver gene mutations and copy number alterations per case is depicted in Fig 3A. These comprise both oncogenes and tumour suppressor genes activated or repressed via different mechanisms. Occasionally different types of events are selected for in the same gene, such as KRAS and ERBB2 which both harbour activating mutations and amplifications in 19% and 18% of cases respectively. Passenger mutations occur by chance in most driver genes. To quantify this we have used the observed:expected mutation ratios (calculated by dNdScv) to estimate the percentage of driver mutations in each gene and in different mutation classes. For many genes, only specific mutation classes appear to be under selection. Many tumour suppressor genes; ARID2, RNF43, ARID1B for example, are only under selection for truncating mutations; ie splice site, nonsense and frameshift Indel mutations, but not missense mutations which are passengers. However, oncogenes, like ERBB2, only contain missense drivers which form clusters to activate gene function in a specific manner. Where a mutation class is <100% driver mutations, mutational clustering can help us define the driver vs passenger status of a mutation (supplementary figure 6). Clusters of mutations occurring in EAC or mutations on amino acids which are mutation hotspots in other cancer types^32^ (supplementary table 5) are indicated in Fig 3A. Novel EAC drivers of particular interest include B2M, a core component of the MHC class I complex and resistance marker for Immunotherapy^33^, MUC6 a secreted glycoprotein involved in gastric acid resistance and ABCB1 a channel pump protein which is associated with multiple instances of drug resistance^34^. We note that several of these drivers have been previously associated with gastric and colorectal cancer (supplementary table 6)^14,35^. Lollipop plots showing primary sequence distribution of mutations in these genes are provided (supplementary data).

**Figure 3.**
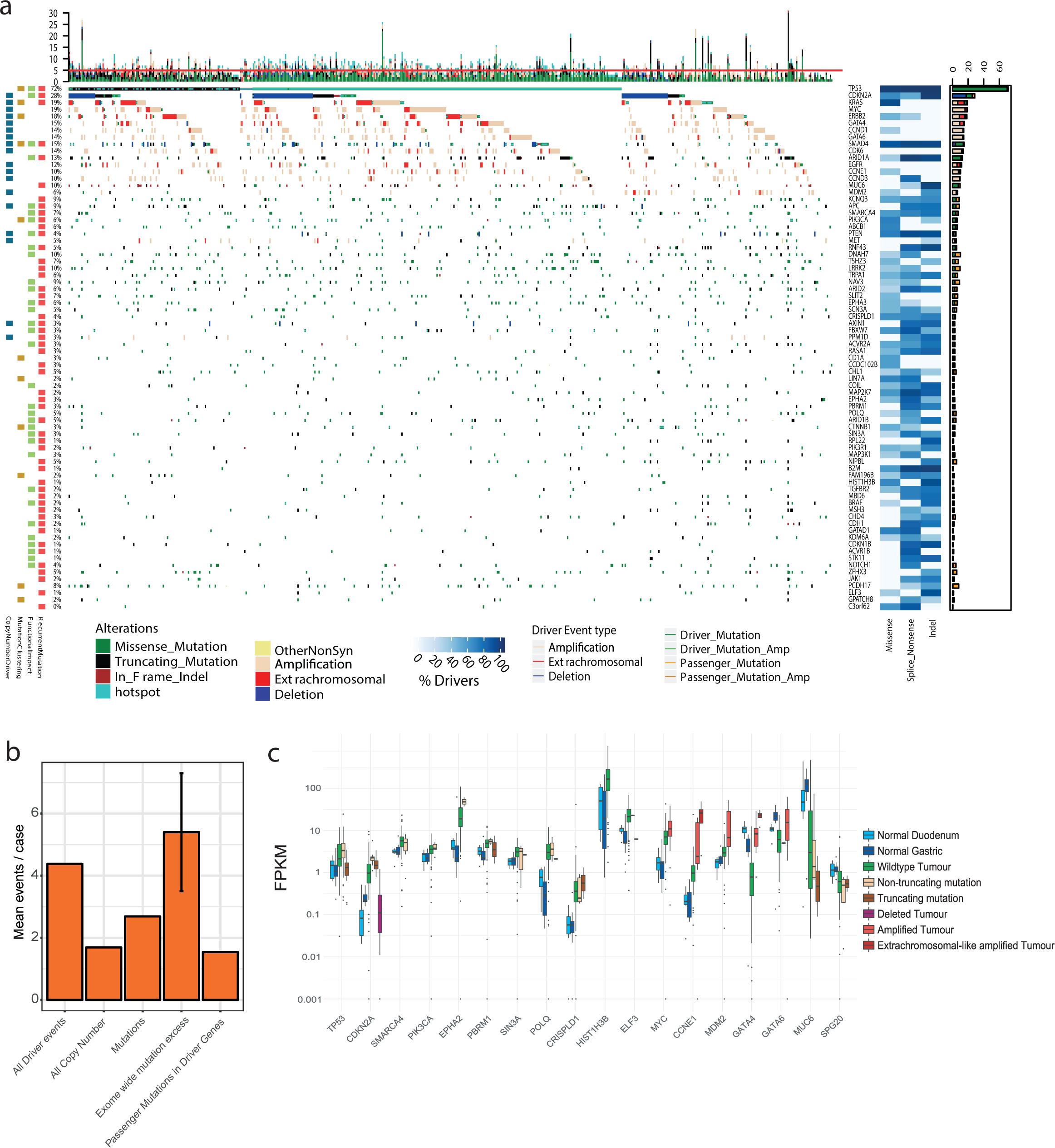
The driver gene landscape of Esophageal Adenocarcinoma. **a**. Driver mutations or CNVs are shown for each patient. Amplification is defined as >2 Copy number adjusted ploidy (2 x ploidy of that case) and extrachromosomal amplification as >10 Copy number adjusted ploidy (10 x ploidy for that case). Driver associated features for each driver gene are displayed to the left. On the right the percentages of different mutation and copy number changes are displayed, differentiating between driver and passenger mutations using dNdScv, and the % of predicted drivers by mutation type is shown. Above the plot are the number of driver mutations per sample with an indication of the mean (red line = 5). **b**. Assessment of driver event types per case and comparison to exome-wide excess of mutations generated by dNdScv. **c**. Expression changes in EAC driver genes in comparison to normal intestinal tissues. Genes with expression changes of note are shown.

The identification of driver events provides a rich information about the molecular history of each EAC tumour. We detect a median of five events in driver genes per tumour (IQR = 3-7, Mean = 5.6) and only a very small fraction of cases have no such events detected (6 cases, 1%). When we remove the predicted percentage of passenger mutations using dnds ratios we find a mean of 4.4 true driver events per case which derive more commonly from mutations than CN events (Fig 3B). Using hierarchal clustering of drivers we noted that TP53 mutant cases had significantly more CN drivers (Wilcox test, p = 0.0032, supplementary figure 7). dNdScv, one of the driver gene detection methods used, also analyses the genome-wide excess of non-synonymous mutations based on expected mutation rates to assess the total number of driver mutations across the exome which is calculated at 5.4 (95% CIs: 3.5-7.3) in comparison to 2.7 driver mutations which we calculate in our gene-centric analysis after passenger removal. This suggests low frequency driver genes may be prevalent in the EAC mutational landscape (see discussion). Further analysis suggests these missing mutations are mostly missense mutations and our gene-centric analysis captures almost all predicted splice and nonsense drivers (supplementary figure 8). Some of our methods use enrichment of nonsense and splice mutations as a marker of driver genes and hence have a higher sensitivity for these mutations.

To better understand the functional impact of driver mutations we analysed expression of driver genes with different mutation types and compared their expression to normal tissue RNA, which was sequenced alongside our tumour samples (Fig 3C). Since surrounding squamous epithelium is a fundamentally different tissue, from which EAC does not directly arise, we have used duodenum and gastric cardia samples as gastrointestinal phenotype controls, likely to be similar to the, as yet unconfirmed, tissue of origin in EAC. A large number of driver genes have upregulated expression in comparison to normal controls, for example TP53 has upregulated RNA expression in WT tumour tissue and in cases with missense (see non-truncating Fig 3C) mutations but RNA expression is lost upon gene truncation. In depth analysis of different TP53 mutation types reveals significant heterogeneity within non-truncating mutations, for example R175H mutations correlate with low RNA expression (supplementary figure 9). Normal tissue expression of CDKN2A suggests that CDKN2A is generally activated in EAC, likely due to genotoxic or other cancer-associated stresses^36^ and returns to physiologically normal levels when deleted. Heterogeneous expression in WT CDKN2A cases suggest a different mechanism of inhibition such as methylation in some cases. Overexpression of other genes in wild type tumours, such as SIN3A, may confer a selective advantage due to their oncogenic properties, in this case cooperating with MYC, which is also overexpressed in EACs (Fig 3C). A smaller number of driver genes are downregulated in EAC tissue-3/4 of these (GATA4, GATA6 and MUC6) are involved in the differentiated phenotype of gastrointestinal tissues and may be lost with tumour de-differentiation. Driving alterations in these genes have been observed in other GI cancers^14,37,38^ however their oncogenic mechanism is unknown. In most genes we did not observe expression loss at the RNA level with truncation, for instance ARID1A (supplementary figure 10).

### Dysregulation of specific pathways and processes in EAC

It is known that selection preferentially dysregulates certain functionally related groups of genes and biological pathways in cancer^39^. This phenomenon is highly evident in EAC, as shown in Fig 4 which depicts the functional relationships between EAC drivers. This provides greater functional homogeneity to the landscape of driver events.

**Figure 4.**
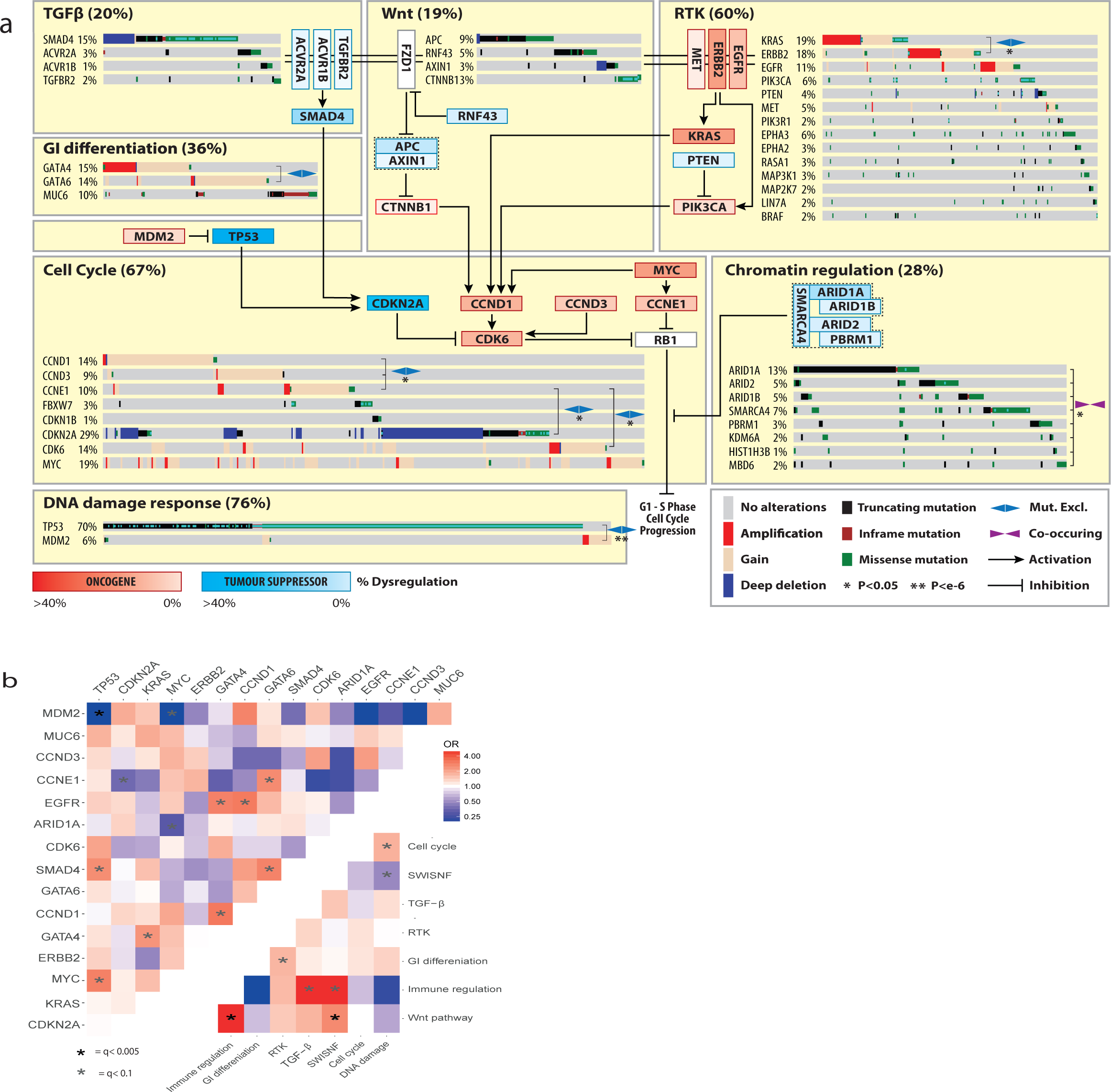
Biological pathways undergoing selective dysregulation in EAC. **a**. Biological Pathways dysregulated by driver gene mutation and/or CNVs. WT cases for a pathway are not shown. Inter and intra-pathway interactions are described and mutual exclusivities and/or associations between genes in a pathway are annotated. GATA4/6 amplifications have a mutually exclusive relationship although this does not reach statistical significance (fisher’s exact test p=0.07 OR =0.52). **b**. Pairwise assessment of mutual exclusivity and association in EAC driver genes and pathways.

While TP53 is the dominant driver in EAC, 28% of cases remain TP53 wildtype. MDM2 is a E3 ubiquitin ligase that targets TP53 for degradation. Its selective amplification and overexpression is mutually exclusive with TP53 mutation suggesting it can functionally substitute the effect of TP53 mutation via its degradation. Similar mutually exclusive relationships are observed between; KRAS and ERBB2, GATA4 and GATA6 and Cyclin genes (CCNE1, CCND1 and CCND3). Activation of the Wnt pathway occurs in 19% of cases either by mutation of phospho-residues at the N terminus of β-catenin, which prevent degradation, or loss of Wnt destruction complex components like APC. Many different chromatin modifying genes, often belonging to the SWI/SNF complex, are also selectively mutated (31% of cases). In contrast SWI/SNF genes are co-mutated significantly more often than we would expect by chance (fisher’s exact test, p<0.01 see methods), suggesting an increased advantage to further mutations once one has been acquired. We also assessed mutual exclusivity and co-occurrence in genes in different pathways and between pathways themselves (Fig 4B). Of particular note are co-occurring relationships between TP53 and MYC, GATA6 and SMAD4, Wnt and Immune pathways as well as mutually exclusive relationships between ARID1A and MYC, gastrointestinal (GI) differentiation and RTK pathways and SWI-SNF and DNA-Damage response pathways. Wnt dysregulation has been previously linked to immune escape^40^ and interestingly was also associated with hyper-mutated cases (> 50,000 SNVs or Indels, fisher’s exact test, p = 0.021, OR= 2.4). We were able to confirm some of these relationships in independent cohorts in different cancer types (supplementary table 7) suggesting some of these may be pan-cancer phenomenon. As shown in Fig 4, all of these pathways interact to stimulate the G1 to S phase transition of the cell cycle via promoting phosphorylation of Rb, although many of these pathways have multiple oncogenic or tumour suppressive functions.

A number of other driver genes have highly related functional roles including core transcriptional components (TAF1 and POLQ), drivers of immune escape (JAK1 and B2M^33^), cell adhesion receptors (CDH1, CHDL and PCDH17), core ribosome components (ELF3 and RPL22), core RNA processing components (GPATCH8 and COIL), ion channels (KCNQ3 and TRPA1) and Ephrin type-A receptors (EPHA2 and EPHA3).

### Clinical significance of driver variants

Events undergoing selection during cancer evolution influence tumour biology and thus impact tumour aggressiveness, response to treatment and patient prognosis as well as other clinical parameters. Clinical-genomic correlations can provide useful biomarkers but also give insights into the biology of these events.

Univariate Cox regression was performed for events in each driver gene with driver events occurring in greater than 5% of EACs (ie after removal of predicted passengers, 16 genes) to detect prognostic biomarkers (Fig 5A). Events in two genes conferred significantly poorer prognosis after multiple hypothesis correction, GATA4 amplification (HR: 0.54, 95% CI: 0.38 – 0.78, *P* value = 0.0008) and SMAD4 mutation or homozygous deletion (HR: 0.60, 95% CI: 0.42 – 0.84, *P* value = 0.003). Both genes remained significant in multivariate Cox regression including pathological TNM staging, resection margin, curative vs palliative treatment intent and differentiation status (GATA4 = HR adjusted: 0.47, 95% CIs adjusted: 0.29 - 0.76, *P* value = 0.002 and SMAD4 = HR adjusted: 0.61, 95% CI adjusted: 0.40 – 0.94, *P* value = 0.026). 31% of EACs contain either SMAD4 mutation or homozygous deletion or GATA4 amplification and cases with both genes altered had a poorer prognosis (Fig 5B). We validated the poor prognostic impact of SMAD4 events in an independent TCGA gastroesophageal cohort (HR = 0.58, 95% CI = 0.37 – 0.90, *P* value =0.014) (Fig 5C) and we also found GATA4 amplifications were prognostic in a cohort of TCGA pancreatic cancers (HR = 0.38 95% CI: 0.18 – 0.80, *P* value = 0.011) (Fig 5D), the only available cohort containing a feasible number of GATA4 amplifications. The prognostic impact of GATA4 has been suggested in previously published independent EAC cohort^17^ although it did not reach statistical significance after FDR correction and SMAD4 expression loss has been previously linked to poor prognosis in EAC^41^. We also noted stark survival differences between cases with SMAD4 events and cases in which TGFβ receptors were mutated (Fig 5E, HR = 5.6, 95% CI: 1.7 – 18.2, *P* value = 0.005) in keeping with the biology of the TGFβ pathway where non-SMAD TGFβ signalling is known to be oncogenic^42^.

**Figure 5.**
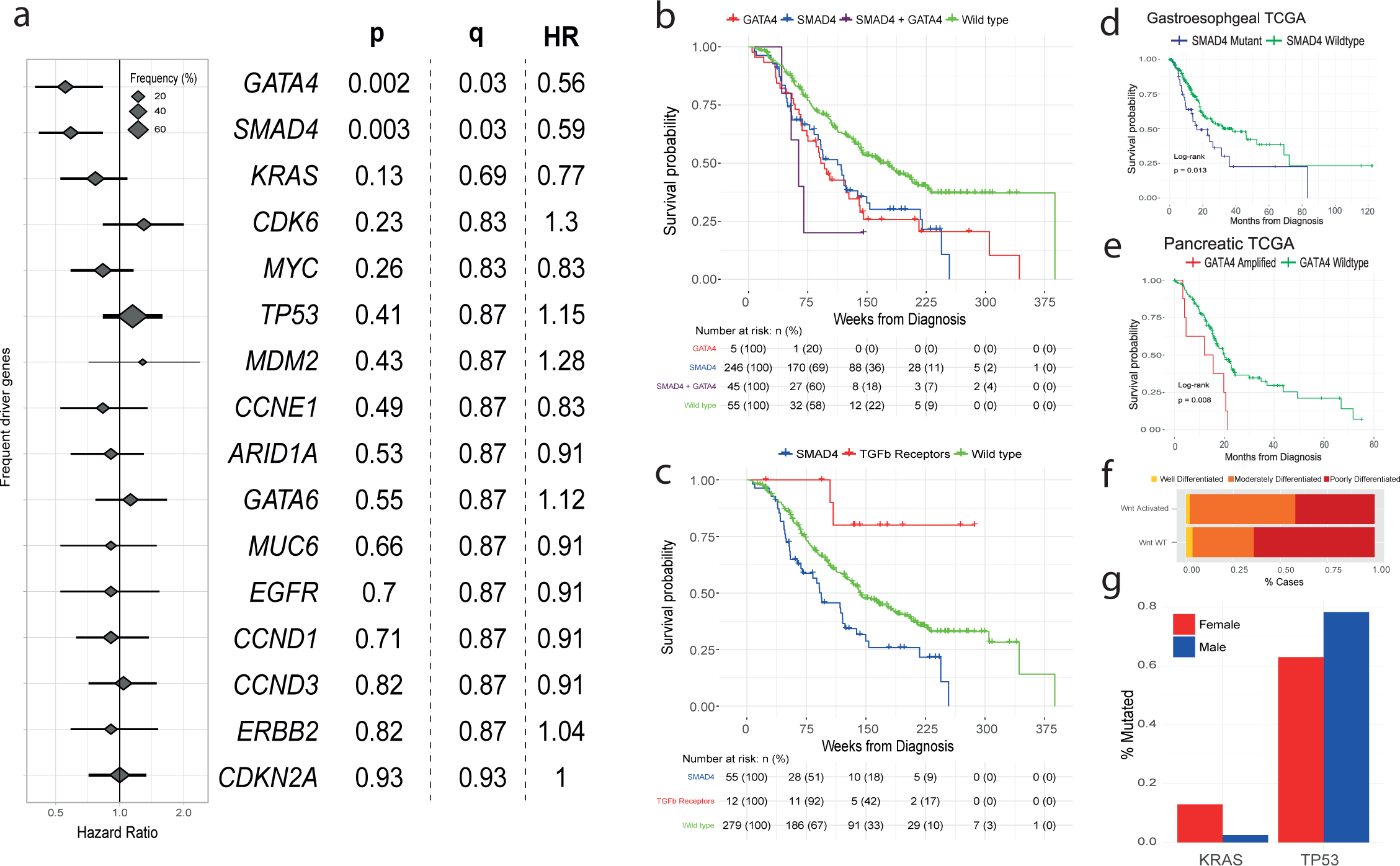
Clinical significance of Driver events in EAC. **a**. Hazard rations and 95% confidence intervals for Cox regression analysis across all drivers genes with at least a 5% frequency of driver alterations * = q < 0.05 after BH adjustment. **b**. Kaplan-Meier curves for EACs with different status of significant prognostic indicators (GATA4 and SMAD4). **c**. Kaplan-Meier curves for different alterations in the TGFbeta pathway. **d**. Kaplan-Meier curves showing verification GATA4 prognostic value in GI cancers using a pancreatic TCGA cohort. **e**. Kaplan-Meier curves showing verification SMAD4 prognostic value in Gastroesophageal cancers using a gastroesophageal TCGA cohort. **f**. Differentiation bias in tumours containing events in Wnt pathway driver genes. **g**. Relative frequency of KRAS mutations and TP53 mutations driver gene events in females vs males (fishers exact test).

In additional to survival analyses we also assessed driver gene events for correlation with various other clinical factors including differentiation status, sex, age and treatment response. We found Wnt pathway mutations had a strong association with well differentiated tumours (p=0.001, OR = 2.9, fisher’s test, see methods, Fig 5F). We noted interesting differences between female (n=81) and male (n=470) cases. Female cases were enriched for KRAS mutation (p = 0.001, fisher’s exact test) and TP53 wildtype status (p = 0.006, fisher’s exact test) (Fig 5G). This is of particular interest given the male predominance of EAC^3^.

### Targeted therapeutics using EAC driver events

The biological distinctions between normal and cancer cells provided by driver events can be used to derive clinical strategies for selective cancer cell killing. To investigate whether the driver events in particular genes and/or pathways might sensitise EAC cells to certain targeted therapeutic agents we used the Cancer Biomarkers database^43^. We calculated the percentage of our cases which contain EAC-driver biomarkers of response to each drug class in the database (summary shown Fig 6A, and full data supplementary table 8). Aside from TP53, which has been problematic to target clinically so far, we found a number of drugs with predicted sensitivity in >10% of EACs including EZH2 inhibitors for SWI/SNF mutant cancers (23%, and 33% including other SWI/SNF EAC drivers), and BET inhibitors which target KRAS activated and MYC amplified cases (25%). However, by far the most significantly effective drug was predicted to be CDK4/6 inhibitors where >50% of cases harboured sensitivity causing events in the receptor tyrosine kinase (RTK) and core cell cycle pathways (eg in CCND1, CCND3 and KRAS).

**Figure 6.**
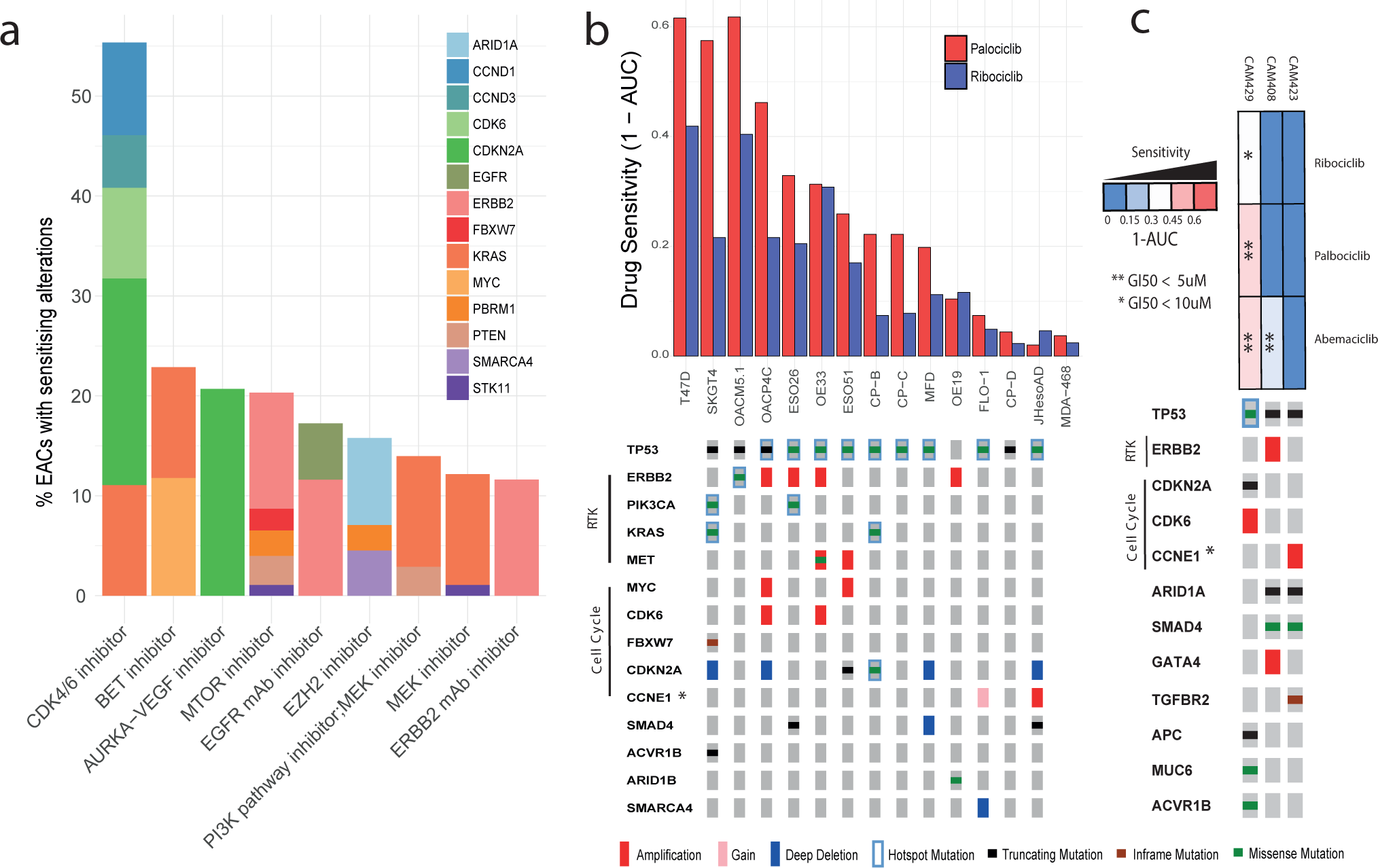
CDK4/6 inhibitors in EAC. **a**. Drug classes for which sensitivity is indicated by EAC driver genes with data from the Cancer Biomarkers database^36^. **b**. Area under the curve (AUC) of sensitivity is shown in a panel of 13 EAC and Be high grade dysplasia cell lines with associated WGS and their corresponding driver events, based on primary tumour analysis. Also AUC is shown for two control lines T47D, an ER +ve breast cancer line (+ve control) and MDA-MB-468 a Rb negative breast cancer (-ve control). *CCNE1 is a known marker of resistance to CDK4/6 inhibitors due to its regulation of Rb downstream of CDK4/6 hence bypassing the need for CDK4/6 activity (see figure 4). c. Response of organoid cultures to three FDA approved CDK4/6 inhibitors and corresponding driver events.

To verify that these driver events would also sensitise EAC tumours to such inhibitors we used a panel of thirteen EAC or Barrett’s HGD cell lines, which share similar genomic changes and driver events^44,45^, which have undergone whole genome sequencing^46^ and assessed them for presence of EAC driver events (Fig 6B). The mutational landscape of these lines was broadly representative of EAC tumours. We found that the presence of cell cycle and or RTK activating driver events was highly correlated with response to two FDA approved CDK4/6 inhibitors, Ribociclib and Palbociclib and several cell lines were sensitive below maximum tolerated blood concentrations in humans (Fig 6B, supplementary table 9, supplementary figure 11)^47^. Such EAC cell lines had comparable sensitivity to T47D which is derived from an ER +ve breast cancer where CDK4/6 inhibitors have been FDA approved. We noted three cell lines without sensitising events which were highly resistant, with little drug effect even at 4000 nanomolar concentrations, similar to a known Rb mutant resistant line breast cancer cell line (MDA-MB-468). Two of these three cell lines harbour amplification of CCNE1 which is known to drive resistance to CDK4/6 inhibitors by bypassing CDK4/6 and causing Rb phosphorylation via CDK2 activation^48^. To verify these effects in a more representative model of EAC we treated three whole genome sequenced EAC organoid cultures^49^ with Palbociclib and Ribociclib as well as a more recently approved CDK4/6 inhibitor, Abemaciclib. As was observed in cell lines, Cell cycle and RTK driver events were present only in the more sensitive organoids and CCNE1 activation in the most resistant (Fig 6C). We found Abemaciclib to be significantly more potent in comparison to both other CDK4/6 inhibitors, both in organoids and cell lines (supplementary figure 10). We note that the maximum tolerated blood doses of Abermaciclib achieved in the clinic were also higher than the other CDK4/6 inhibitors^50^, within the range of sensitivity achieved in several cell lines and organoids cultures.

## Discussion

We present here a detailed catalogue of coding and non-coding genomic events that have been selected for during the evolution of esophageal adenocarcinoma. These events have been characterised in terms of their relative impact, related functions, mutual exclusivity and co-occurrence and expression in comparison to normal tissues, producing insights into EAC biology. We have used this set of biologically important gene alterations to identify prognostic biomarkers and actionable genomic events for personalised medicine.

While clinical annotation and matched RNA data is a strength of this study, in some cases we may have been unable to assess selected variants for survival associations or expression changes which were detected in the full 551 cohort, due to lack of representation in clinically annotated or RNA matched sub cohorts. Despite rigorous analyses to detect selected events, assessment of the global excess of mutations by dNdScv suggests we are unable to detect all events selected in EAC, similar to many other cancer types^22^. All driver gene detection methods which we have used rely on driver mutation re-occurrence in a gene to some degree. Many of these undetected driver mutations are hence likely to be spread across a large number of genes whereby each is mutated at low frequency across EAC patients. This tendency for low frequency EAC drivers may be responsible for the low yield of MutsigCV in previous cohorts and may suggests that C-type cancers such as EAC, are not less ‘mutation-driven’ than M-type cancers but rather that their mutational drivers are spread across a larger number of genes^5^. The identification of these very low frequency mutations will require substantially different detection techniques to those which are currently in wide spread use and such methods are in development^51^ although they require validation. Undoubtedly many copy number drivers are also left undiscovered and validation of candidates identified here is an important avenue of future work.

While a number of previous reports have attempted to detect EAC drivers, they have had a limited yield per case for a variety of reasons. The first such study^20^ used methods which, despite being well regarded at the time, were subsequently discredited^9^. Hence a number of known false positive genes (EYS, SYNE1 and CNTTAP5) were erroneously reported as drivers, along with an additional unknown number of genes. Since then a number of reports, including our own, on medium and large cohort sizes using MutsigCV^10,11,18^ were only able to detect a small number of mutational driver genes (7, 5 and 15 in each study). By using both a large cohort and more comprehensive methodologies we have significantly increased this figure to 66 mutational driver genes (excluding CN drivers). Detection of driver CNAs has previously relied on GISTIC to detect recurrently mutated regions^10,15-18^ but no analyses have been performed to evidence which genes in these large regions are true drivers. Many of the genes annotated by such papers are unlikely to be CN drivers from this analysis due to their lack of expression modulation with CNAs (eg YEATS4 and MCL1), the role of recurrent heterozygous losses to drive LOH in some mutational drivers (ARID1A and CDH11) or their association with fragile sites (PDE4D, WWOX, FHIT). Conversely, we have been able to identify novel EAC copy number drivers (eg CCND3, AXIN1, PPM1D and APC).

A number of discoveries made in this work require further investigation. Functional characterisation of many of the driver genes described is needed to understand why they are advantageous to EAC tumours and how they modify EAC biology. Particularly interesting are the GI specific genes GATA4, GATA6 and MUC6 which modulate prognosis and have expression loss during the transition from normal to tumour tissue. Biological pathways and processes that are selectively dysregulated deserve particular attention in this regard as do the gene pairs or groups with mutually exclusive or co-occurring relationships such as MYC and TP53 or SWI/SNF factors, suggestive of particular functional relationships. Prospective clinical work to verify and implement SMAD4 and GATA4 biomarkers in this study would be worthwhile. While EAC is a poor prognosis cancer type, significant heterogeneity of survival outcome makes triaging patients in treatment groups an important part of clinic practice which could be improve using better prognostication. Whole genome or whole exome sequencing may be impractical for use in the clinic, however targeted NGS panels to detect mutations and copy number alterations have been implemented to detect genomic biomarkers in a cost effective and sensitive manner for some cancer types^52^. In EAC development of a customised panel is likely to be required on the basis of this analysis. A number of targeted therapeutics may provide clinic benefit to EAC cases based on their individual genomic profile. In particular CDK4/6 inhibitors deserve considerable attention as an option for EAC treatment as they are, by a significant margin, the treatment to which the most EACs harbour sensitivity-causing driver events, excluding TP53 as an unlikely therapeutic biomarker. The in vitro validation of these biomarkers for CDK4/6 inhibitors in EAC is also persuasive of possible clinical benefit using a targeted approach.

In summary this work provides a detailed compendium of mutations and copy number alterations undergoing selection in EAC which have functional and clinical impact on tumour behaviour. This comprehensive study provides us with useful insights into the nature of EAC tumours and should pave the way for evidence based clinical trials in this poor prognosis disease.

## Author contributions

RCF and AMF conceived the overall study. AMF and SJ analysed the genomic data and performed statistical analyses. RCF, AMF and XL designed the experiments. AMF, XL and JM performed the experiments. GC contributed to the Structural variant analysis and data visualisation. SK helped compile the clinical data and aided statistical analyses. JP and SA produced and QC’ed the RNA-seq data. EO aided the whole genome sequencing of EAC cell lines. SM and NG coordinated the clinical centres and were responsible for sample collections. ME benchmarked our mutation calling pipelines. MO led the pathological sample QC for sequencing. LB and GD ran variant calling pipelines. RCF and ST supervised the research. RCF and ST obtained funding. AMF and RCF wrote the manuscript. All authors approved the manuscript.

The authors declare no competing interests.

Sequencing data will be deposited in a publicly assessable database before publication Code associated with the analysis is available upon request.

The study was registered (UKCRNID 8880), approved by the Institutional Ethics Committees (REC 07/H0305/52 and 10/H0305/1), and all subjects gave individual informed consent.

OCCAMS was funded by a programme grant from Cancer Research UK (RG66287). We thank the Human Research Tissue Bank, which is supported by the National Institute for Health Research (NIHR) Cambridge Biomedical Research Centre, from Addenbrooke’s Hospital. Additional infrastructure support was provided from the CRUK funded Experimental Cancer Medicine Centre.

## Acknowledgements

We would like to thank Dr. Adam Bass and Dr. Nic Waddel for providing data in ^Dulak et al 2013^ and Nones et al 2014 respectively, also included in our previous publication Secrier et al 2016. Inclusion of this data allowed augmentation of our ICGC cohort and a greater sensitivity for the detection of EAC driver genes.

## Supplementary figure legends

**Supplementary figure 1.** Distribution of small scale mutations (SNVs and Indels) across the 551 EAC cohort. Red line indicates the median mutations per case (6.4)

**Supplementary Figure 2. Concordance between driver gene detection methods. A.** Hierarchical clustering between tools based on gene identified. **B** Genes identified by each tool.

**Supplementary Figure 3.** Frequency and significance of EAC non-coding drivers from ActiveDriverWGS. **a.** The observed and expected mutation counts found on each element in ActiveDriverWGS. **b.** The fdr for each element in ActiveDriverWGS.

**Supplementary Figure 4.** Frequency of Extrachromosomal like events (CN adjusted Ploidy >10) in GISTIC amplification peaks and presence of high confidence drivers in those peaks indicated.

**Supplementary figure 5. Examples of Normal amplification (PLiody-adjusted CN >2 & <10) and Extrachromosomal-like amplification (ploidy-adjusted CN >10) events**. 1-10 = Extrachromosomal-like amplification and 11-20 = Normal amplification events. Events were picked at random using runif() function in R. SV and CNAs surrounding events are shown. Features indicative of extrachromosomal double minute (DM) formations include sharp, large CN steps, SVs with high read support at the edges of these steps and when not derived from a continuous region of the genome CN regions in the DM may have the same CN status (taking into account other additional events which may have occurred in that region). These features are enriched in the extrachromosomal-like events, although example 20 may be a low-copy number extrachromosomal event. It should be noted that SV calling using short read sequencing techniques such as in this study has a relatively low sensitivity and accuracy for the precise localisation of many SV break points. Examples continue over four pages.

**Supplementary Figure 6**. A scheme demonstrating how to use mutational clustering along with dnds ratios to estimate the probability of a particular mutation being a driver. In this case the dnds ratio suggests 2/3 of missense mutations are drivers hence 10/15. 8 missense mutation lie in a mutational cluster, in this case of known significance in the N-terminal of B-Catenin, making it likely that these are drivers and hence most (2/7) other mutations are passengers. Similarly, mutations on amino acids known to be hyper mutated in other cancer types (see Supplementary table 5, for instance if we found a single KRAS G12 mutation) can be considered likely drivers.

**Supplementary Figure 7. Hierarchical Clustering of samples based on presence of driver variants with genes ordered by pathway membership.**

**Supplementary Figure 8.** A detailed breakdown of mutation and copy number types per case and a breakdown of exome wide dnds excess for different mutation types (note that exome wide indel cannot be calculated excess as they have no synonymous mutation equivalent, although a null model is used in the per gene dnds method to use them to detect driver genes). Error bars indicate 95% confidence intervals for exome-wide dnds mutation excess assessment.

**Supplementary Figure 9.** TP53 expression in different TP53 mutation types in comparison to TP53 WT tumours and normal duodenum and gastric cardia tissues.

**Supplementary Figure 10.** Expression of all EAC driver genes across different genomic states for the gene in question in 116 EAC tumours, and in comparison to duodenum and gastric cardia tissues.

**Supplementary Figure 11.** Growth inhibition responses of EAC cell lines and control lines to CDK4/6 inhibitors Palbociclib and Ribociclib. A subset of cell lines also received treatment with Abemaciclib which shows efficacy in such cell lines as well as in organoids (Fig 6C).

## Methods

### Cohort, sequencing and calling of genomic events

380 cases (69%) of our EAC cohort were derived from the esophageal adenocarcinoma WGS ICGC study, for which samples are collected through the UK wide OCCAMS (Oesophageal Cancer Classification and Molecular Stratification) consortium. The procedures for obtaining the samples, quality control processes, extractions and whole genome sequencing are as previously described^18^. Strict pathology consensus review was observed for these samples with a 70% cellularity requirement before inclusion. Comprehensive clinical information was available for the ICGC-OCCAMS cases. In addition, previously published samples were included in the analysis from Dulak et al 2013^20^ – 139 WES and 10 WGS (total 27%) and Nones et al 2014^21^ with 22 WGS samples (4%) to total 551 genome characterised EACs. RNA-seq data was available from our ICGC WGS samples (116/380). BAM files for all samples (include those from ^Dulak et al 2013^ and Nones et al 2014) were run through our alignment (BWA-MEM), mutation (Strelka), copy number (ASCAT) and structural variant (Manta) calling pipelines, as previously described^18^. Our methods were benchmarked against various other available methods and have among the best sensitivity and specificity for variant calling (ICGC benchmarking excerise^53^). Mutation and copy number calling on cell lines was performed as previously described^46^.

Total RNA was extracted using All Prep DNA/RNA kit from Qiagen and the quality was checked on Agilent 2100 Bioanalyzer using RNA 6000 nano kit (Agilent). Qubit High sensitivity RNA assay kit from thermo fisher was used for quantification. Libraries were prepared from 250ng RNA, using TruSeq Stranded Total RNA Library Prep Gold (Ribo-zero) kit and ribosomal RNA (nuclear, cytoplasmic and mitochondrial rRNA) was depleted, whereby biotinylated probes selectively bind to ribosomal RNA molecules forming probe-rRNA hybrids. These hybrids were pulled down using magnetic beads and rRNA depleted total RNA was reverse transcribed. The libraries were prepared according to Illumina protocol^54^. Paired end 75bp sequencing on HiSeq4000 generated the paired end reads. For normal expression controls we chose gastric cardia tissue, from which some hypothesise Barrett’s may arise, and duodenum which contains intestinal histology, including goblet cells, which mimics that of Barrett’s. We did not use Barrett’s tissue itself as a normal control given the heterogeneous and plentiful phenotypic and genomic changes which it undergoes early in its pathogenesis.

### Analysing EAC mutations for selection

To detect positively selected mutations in our EAC cohort, a multi-tool approach across various selection related ‘Features’ (Recurrance, Functional impact, Clustering) was implemented in order to provide a comprehensive analysis. This is broadly similar to several previous approaches^8,12^. dNdScv^22^, MutsigCV^9^, e-Driver^26^, ActivedriverWGS and e-Driver3D^27^ were run using the default parameters. To run OncodriverFM^24^, Polyphen^55^ and SIFT^56^ were used to score the functional impact of each missense non-synonomous mutation (from 0, non-impactful to 1 highly impactful), synonymous mutation were given a score of 0 impact and truncating mutations (Non-sense and frameshift mutations) were given a score of 1. Any gene with less than 7 mutations, unlikely to contain detectable drivers using this method, was not considered to decrease the false discovery rate. OncodriveClust was run using a minimum cluster distance of 3, minimum number of mutations for a gene to be considered of 7 and with a stringent probability cut off to find cluster seeds of p = Ex10^−13^ to prevent infiltration of large numbers of, likely, false positive genes. For all tool outputs we undertook quality control including Q-Q plots to ensure no tool produces inflated q-values and each tool produced at least 30% known cancer genes. Two tools were removed from the analysis due to failure for both of these parameters at quality control (Activedriver^57^ and Hotspot^32^). For three of the QC-approved tools (dNdScv, OncodriveFM, MutsigCV) where this was possible we also undertook an additional fdr reducing analysis by re-calculating q values based on analysis of known cancer genes only^22,28,29^ as has been previously implemented^22,58^. Significance cut offs were set at q<0.1 for coding genes. Tool outputs were then put through various filters to remove any further possible false positive genes. Specifically, genes where <50% of EAC cases had no expression (TPM<0.1) in our matched RNA-seq cohort were removed and, using dNdScv, genes with no significant mutation excess (observed: expected ratio > 1.5:1) of any single mutation type were also removed. We also removed two (MT-MD2, MT-MD4) mitochondrial genes which were highly enriched for truncating mutations and were frequently called in OncodriveFM as well as other tools. This is may be due to the different mutational dynamics, caused by ROS from the mitochondrial electron transport chain, and the high number of mitochondrial genomes per cell which enables significantly more heterogeneity. These factors prevent the tools used from calculating an accurate null model for these genes however they may be worthy of functional investigation. For non-coding elements called by ActivedriverWGS filtering for expression or dnds was not possible and dispite recent benchmarking^30^ are not so well established. Hence we took a more cautious approach with general significance cut offs of q < 0.001 and q < 0.1 for previously identified elements in PCAWG^12^. Q values were not recalculated for Driver elements only but q < 0.1 for known elements was based on all elements. To calculate exome-wide mutational excess hypermutated cases (>500 exonic mutations) were removed and the global non-synonymous dnds ratios were applied to all dndscv annotated mutations excluding “synonymous” and “no SNV” annotations as described in Martincorena et al^22^.

### Detecting selection in CNVs

ASCAT raw CN values were used to detected frequently deleted or amplified regions of the genome using GISTIC2.0^15^. To determine which genes in these regions confer a selective advantage, CNVs from each gene within a GISTIC identified loci were correlated with TPM from matched RNA-seq in a sub-cohort of 116 samples and with mutations across all 551 samples. To call copy number in genes which spanned multiple copy number segments in ASCAT we considered the total number of full copies of the gene (ie the lowest total copy number). Occasionally ASCAT is unable to confidently call the copy number in a highly aberrant genomic regions. We found that the expression of genes in such regions matched well what we would expect given the surrounding copy number and hence we used the mean of the two adjacent copy number fragments to call copy number in the gene in question. We found amplification peak regions identified by GISTIC2.0 varied significantly in precise location both in analysis of different sub-cohorts and when comparing to published GISTIC data from EACs^10,16,17^. A peak would often sit next to but not overlapping a well characterised oncogene or tumour suppressor. To account for this, we widened the amplification peak sizes upstream and downstream by twice the size of each peak to ensure we captured all possible drivers. Our expression analysis allows us to then remove false positives from this wider region and called drivers were still highly enriched for genes closer to the centre of GISTIC peak regions.

To detect genes in which amplification correlated with increased expression we compared expression of samples with a high CN for that gene (above 10^th^ percentile CN/Ploidy) with those which have a normal CN (median +/− 1) using the Wilcox rank-sum test and using the specific alternative hypothesis that high CN would lead to increased expression. Q-values were then generated based on Benjamini & Hochberg method, not considering genes without significant expression in amplified samples (at least 75% amplified samples with TPM > 0.1) and considering q<0.001 as significant. We also included an additional known driver gene only FDR reduction analysis as previously described for mutational drivers with q<0.1 considered as significant given the additional evidence for these genes in other cancer types. We also included MYC despite its q= 0.11 for expression correlation. This is due to frequent non-amplification associated overexpression of MYC when compared to normal controls and otherwise MYC is well evidence by a very close proximity to the peak centre (top 4 genes) and its high rate of amplification (19%). We took the same approach to detect genes in which homozygous deletion correlated with expression loss. Expression modulation was a highly specific marker for known CN driver genes and was not a widespread feature in most recurrently copy number variant genes. However, while expression modulation is a requirement for selection of CNV only drivers, it is not sufficient evidence alone and hence we grouped such genes into those which have been characterised as drivers previously in other cancer types (high confidence EAC CN drivers) and other genes (Candidate EAC CN drivers) which await functional validation. We used fragile site regions detected in Wala et al 2017^59^. We also defined regions which may be recurrently heterozygous deleted, without any significant expression modulations, to allow LOH of tumour suppressor gene mutations. To do this we analysed genes with at least 5 mutations in the matched RNA cohort for association between LOH (ASCAT minor allele = 0) and mutation using fisher’s exact test and generated q values using the Benjamini & Hochberg method. The analysis was repeated on known cancer genes only for reduced FDR and q < 0.05 considered significant for both analyses. For those high confidence drivers we chose to define amplification as CN/ploidy (referred to as Ploidy adjusted copy number) this produces superior correlation with expression. We chose a cut off for amplification at CN/ploidy = 2 as has been previously used, and as causes a highly significant increase in expression in our CN-driver genes.

### Pathways and relative distributions of genomic events

The relative distribution of driver events in each pathway was analysed using a fisher’s exact test in the case of pair-wise comparisons including WT cases. In the case of multi-gene comparisons such as the Cyclins we calculate the p value and odds ratio for each pair in the group by fisher’s exact test and combine p values using the Fisher method, Genes without comparable Odds ratios to the rest of the genes in question were removed. For this analysis we also remove highly mutated cases (>500 exonic mutations, 41/551) as they bias distribution of genes towards co-occurrence. We repeated this analyses across all pairs of driver genes using BH multiple hypothesis correction. We validated these relationships in independent TGCA cohorts of other GI cancers where we could find cohorts with reasonable numbers of the genomic events in question (not possible for GATA4/6 for instance) using the cBioportal web interface tool^60^.

### Correlating genomics with the clinical phenotype

To find genomic markers for prognosis we undertook univariate Cox regression for those driver genes present in >5% of cases (16) along with Benjamini & Hochberg false discovery correction. We considered only these genes to reduce our false discover rate and because other genes were unlikely to impact on clinical practise given their low frequency in EAC. We validated SMAD4, in the TCGA gastroesophageal cohort which had a comparable frequency of these events, but notably is composed mainly of gastric cancers, and GATA4 in the TCGA pancreatic cohort using the cBioportal web interface tool. We also validated these markers as independent predictors of survival both in respect of each other and stage using a multivariate Cox regression in our 551 case cohort. When assessing for genomic correlates with differentiation phenotypes we found only very few cases with well differentiated phenotypes (<5% cases) and hence for statistical analyses we collapse these cases with moderate differentiation to allow a binary fisher’s exact test to compare poorly differentiated with well-moderate differentiated phenotypes.

### Therapeutics

The cancer biomarker database was filtered for drugs linked to biomarkers found in EAC drivers and supplementary table 6 constructed using the cohort frequencies of EAC biomarkers. 10 EAC cell lines (SKGT4, OACP4C, OACM5.1, ESO26, ESO51, OE33, MFD, OE19, Flo-1 and JHesoAD) and 3 BE high grade dysplasia cell lines (CP-B, CP-C and CP-D) with WGS data^46^ were used in proliferation assays to determine drug sensitivity to CDK4/6 inhibitors, Palbociclib (Biovision) and Ribociclib (Selleckchem). Cell lines were grown in their normal growth media (methods table 1). Proliferation was measured using the Incucyte live cell analysis system (Incucyte ZOOM Essen biosciences). Each cell line was plated at a starting confluency of 10% and growth rate measured across 4-7 days depending on basal proliferation rate. For each cell-line drug combination concentrations of 16, 64, 250, 1000 and 4000 nanomolar were used each in 0.3% DMSO and compared to 0.3% DMSO only. Each condition was performed in at least triplicate. The time period of the exponential growth phase in the untreated (0.3% DMSO) condition was used to calculate GI50 and AUC. Accurate GI50s could not be calculated in cases where a cell line had >50% proliferation inhibition even with the highest drug concentration and hence AUC was used to compare cell line sensitivity. T47D had a highly similar GI50 for Palbociclib to that previously calculated in other studies (112 nM vs 127 nM)^61^. Primary organoid cultures were derived from EAC cases included in the OCCAMS/ICGC sequencing study. Detailed organoid culture and derivation method have been previously described (cite nat comms Li et al). Regarding the drug treatment, the seeding density for each line was optimised to ensure cell growth in the logarithmic growth phase. Cells were seeded in complete medium for 24 hours then treated with compounds at a 5-point 4-fold serial dilutions for 6 days or 12 days. Cell viability was assessed using CellTiter-Glo (Promega) after drug incubation.

## References

1. Ferlay J, Soerjomataram I, Dikshit R, et al. Cancer incidence and mortality worldwide: sources, methods and major patterns in GLOBOCAN 2012. Int J Cancer. Mar 1 2015;136(5):E359–386.

2. Coleman HG, Xie SH, Lagergren J. The Epidemiology of Esophageal Adenocarcinoma. Gastroenterology. Jan 2018;154(2):390–405.

3. Smyth EC, Lagergren J, Fitzgerald RC, et al. Oesophageal cancer. Nat Rev Dis Primers. Jul 27 2017;3:17048.

4. Campbell PJ, Getz G, Stuart JM, Korbel JO, Stein LD. Pan-cancer analysis of whole genomes. bioRxiv. 2017.

5. Ciriello G, Miller ML, Aksoy BA, Senbabaoglu Y, Schultz N, Sander C. Emerging landscape of oncogenic signatures across human cancers. Nat Genet. Oct 2013;45(10):1127–1133.

6. Secrier M, Li X, de Silva N, et al. Mutational signatures in esophageal adenocarcinoma define etiologically distinct subgroups with therapeutic relevance. Nat Genet. Oct 2016;48(10):1131–1141.

7. Stratton MR, Futreal PA. Cancer: understanding the target. Nature. Jul 1 2004;430(6995):30.

8. Tamborero D, Gonzalez-Perez A, Perez-Llamas C, et al. Comprehensive identification of mutational cancer driver genes across 12 tumor types. Sci Rep. Oct 2 2013;3:2650.

9. Lawrence MS, Stojanov P, Polak P, et al. Mutational heterogeneity in cancer and the search for new cancer-associated genes. Nature. Jul 11 2013;499(7457):214–218.

10. Integrated genomic characterization of oesophageal carcinoma. Nature. Jan 12 2017;541(7636):169–175.

11. Lin DC, Dinh HQ, Xie JJ, et al. Identification of distinct mutational patterns and new driver genes in oesophageal squamous cell carcinomas and adenocarcinomas. Gut. Aug 31 2017.

12. Rheinbay E, Nielsen MM, Abascal F, et al. Discovery and characterization of coding and non-coding driver mutations in more than 2,500 whole cancer genomes. bioRxiv. 2017.

13. Comprehensive molecular characterization of urothelial bladder carcinoma. Nature. Mar 20 2014;507(7492):315–322.

14. Comprehensive molecular characterization of gastric adenocarcinoma. Nature. Sep 11 2014;513(7517):202–209.

15. Mermel CH, Schumacher SE, Hill B, Meyerson ML, Beroukhim R, Getz G. GISTIC2.0 facilitates sensitive and confident localization of the targets of focal somatic copy-number alteration in human cancers. Genome Biol. 2011;12(4):R41.

16. Dulak AM, Schumacher SE, van Lieshout J, et al. Gastrointestinal adenocarcinomas of the esophagus, stomach, and colon exhibit distinct patterns of genome instability and oncogenesis. Cancer Res. Sep 1 2012;72(17):4383–4393.

17. Frankel A, Armour N, Nancarrow D, et al. Genome-wide analysis of esophageal adenocarcinoma yields specific copy number aberrations that correlate with prognosis. Genes Chromosomes Cancer. Apr 2014;53(4):324–338.

18. Secrier M, Fitzgerald RC. Signatures of Mutational Processes and Associated Risk Factors in Esophageal Squamous Cell Carcinoma: A Geographically Independent Stratification Strategy? Gastroenterology. May 2016;150(5):1080–1083.

19. Zack TI, Schumacher SE, Carter SL, et al. Pan-cancer patterns of somatic copy number alteration. Nat Genet. Oct 2013;45(10):1134–1140.

20. Dulak AM, Stojanov P, Peng S, et al. Exome and whole-genome sequencing of esophageal adenocarcinoma identifies recurrent driver events and mutational complexity. Nat Genet. May 2013;45(5):478–486.

21. Nones K, Waddell N, Wayte N, et al. Genomic catastrophes frequently arise in esophageal adenocarcinoma and drive tumorigenesis. Nat Commun. Oct 29 2014;5:5224.

22. Martincorena I, Raine KM, Gerstung M, et al. Universal Patterns of Selection in Cancer and Somatic Tissues. Cell. Nov 16 2017;171(5):1029–1041 e1021.

23. Wadi L, Uuskula-Reimand L, Isaev K, et al. Candidate cancer driver mutations in super-enhancers and long-range chromatin interaction networks. bioRxiv. 2017.

24. Gonzalez-Perez A, Lopez-Bigas N. Functional impact bias reveals cancer drivers. Nucleic Acids Res. Nov 2012;40(21):e169.

25. Tamborero D, Gonzalez-Perez A, Lopez-Bigas N. OncodriveCLUST: exploiting the positional clustering of somatic mutations to identify cancer genes. Bioinformatics. Sep 15 2013;29(18):2238–2244.

26. Porta-Pardo E, Godzik A. e-Driver: a novel method to identify protein regions driving cancer. Bioinformatics. Nov 1 2014;30(21):3109–3114.

27. Porta-Pardo E, Hrabe T, Godzik A. Cancer3D: understanding cancer mutations through protein structures. Nucleic Acids Res. Jan 2015;43(Database issue):D968–973.

28. Futreal PA, Coin L, Marshall M, et al. A census of human cancer genes. Nat Rev Cancer. Mar 2004;4(3):177–183.

29. Kandoth C, McLellan MD, Vandin F, et al. Mutational landscape and significance across 12 major cancer types. Nature. Oct 17 2013;502(7471):333–339.

30. Shuai S, Gallinger S, Stein LD. DriverPower: Combined burden and functional impact tests for cancer driver discovery. bioRxiv. 2017.

31. Taylor AM, Shih J, Ha G, et al. Genomic and Functional Approaches to Understanding Cancer Aneuploidy. Cancer cell. Apr 9 2018;33(4):676–689 e673.

32. Chang MT, Asthana S, Gao SP, et al. Identifying recurrent mutations in cancer reveals widespread lineage diversity and mutational specificity. Nat Biotechnol. Feb 2016;34(2):155–163.

33. Zaretsky JM, Garcia-Diaz A, Shin DS, et al. Mutations Associated with Acquired Resistance to PD-1 Blockade in Melanoma. N Engl J Med. Sep 1 2016;375(9):819–829.

34. Chen Z, Shi T, Zhang L, et al. Mammalian drug efflux transporters of the ATP binding cassette (ABC) family in multidrug resistance: A review of the past decade. Cancer Lett. Jan 1 2016;370(1):153–164.

35. Giannakis M, Mu XJ, Shukla SA, et al. Genomic Correlates of Immune-Cell Infiltrates in Colorectal Carcinoma. Cell reports. Oct 18 2016;17(4):1206.

36. Pei XH, Xiong Y. Biochemical and cellular mechanisms of mammalian CDK inhibitors: a few unresolved issues. Oncogene. Apr 18 2005;24(17):2787–2795.

37. Comprehensive molecular characterization of human colon and rectal cancer. Nature. Jul 18 2012;487(7407):330–337.

38. Waddell N, Pajic M, Patch AM, et al. Whole genomes redefine the mutational landscape of pancreatic cancer. Nature. Feb 26 2015;518(7540):495–501.

39. Leiserson MD, Vandin F, Wu HT, et al. Pan-cancer network analysis identifies combinations of rare somatic mutations across pathways and protein complexes. Nat Genet. Feb 2015;47(2):106–114.

40. Grasso CS, Giannakis M, Wells DK, et al. Genetic Mechanisms of Immune Evasion in Colorectal Cancer. Cancer discovery. Jun 2018;8(6):730–749.

41. Singhi AD, Foxwell TJ, Nason K, et al. Smad4 loss in esophageal adenocarcinoma is associated with an increased propensity for disease recurrence and poor survival. Am J Surg Pathol. Apr 2015;39(4):487–495.

42. Levy L, Hill CS. Alterations in components of the TGF-beta superfamily signaling pathways in human cancer. Cytokine Growth Factor Rev. Feb-Apr 2006;17(1-2):41–58.

43. Tamborero D, Rubio-Perez C, Deu-Pons J, et al. Cancer Genome Interpreter Annotates The Biological And Clinical Relevance Of Tumor Alterations. bioRxiv. 2017.

44. Weaver JMJ, Ross-Innes CS, Shannon N, et al. Ordering of mutations in preinvasive disease stages of esophageal carcinogenesis. Nature genetics. Aug 2014;46(8):837–843.

45. Ross-Innes CS, Becq J, Warren A, et al. Whole-genome sequencing provides new insights into the clonal architecture of Barrett's esophagus and esophageal adenocarcinoma. Nature genetics. Sep 2015;47(9):1038–1046.

46. Contino G, Eldridge MD, Secrier M, et al. Whole-genome sequencing of nine esophageal adenocarcinoma cell lines. F1000Res. 2016;5:1336.

47. Liston DR, Davis M. Clinically Relevant Concentrations of Anticancer Drugs: A Guide for Nonclinical Studies. Clin Cancer Res. Jul 15 2017;23(14):3489–3498.

48. Herrera-Abreu MT, Palafox M, Asghar U, et al. Early Adaptation and Acquired Resistance to CDK4/6 Inhibition in Estrogen Receptor-Positive Breast Cancer. Cancer Res. Apr 15 2016;76(8):2301–2313.

49. Li X, Francies HE, Secrier M, et al. Organoid cultures recapitulate esophageal adenocarcinoma heterogeneity providing a model for clonality studies and precision therapeutics. Nature communications. Jul 30 2018;9(1):2983.

50. Patnaik A, Rosen LS, Tolaney SM, et al. Efficacy and Safety of Abemaciclib, an Inhibitor of CDK4 and CDK6, for Patients with Breast Cancer, Non-Small Cell Lung Cancer, and Other Solid Tumors. Cancer discovery. Jul 2016;6(7):740–753.

51. D’Antonio M, Ciccarelli FD. Integrated analysis of recurrent properties of cancer genes to identify novel drivers. Genome Biol. May 29 2013;14(5):R52.

52. Zehir A, Benayed R, Shah RH, et al. Mutational landscape of metastatic cancer revealed from prospective clinical sequencing of 10,000 patients. Nat Med. Jun 2017;23(6):703–713.

53. Ding J, McConechy MK, Horlings HM, et al. Systematic analysis of somatic mutations impacting gene expression in 12 tumour types. Nat Commun. Oct 5 2015;6:8554.

54. Nagai K, Kohno K, Chiba M, et al. Differential expression profiles of sense and antisense transcripts between HCV-associated hepatocellular carcinoma and corresponding non-cancerous liver tissue. Int J Oncol. Jun 2012;40(6):1813–1820.

55. Adzhubei I, Jordan DM, Sunyaev SR. Predicting functional effect of human missense mutations using PolyPhen-2. Curr Protoc Hum Genet. Jan 2013;Chapter 7:Unit7 20.

56. Ng PC, Henikoff S. Predicting the effects of amino acid substitutions on protein function. Annu Rev Genomics Hum Genet. 2006;7:61–80.

57. Reimand J, Wagih O, Bader GD. The mutational landscape of phosphorylation signaling in cancer. Sci Rep. Oct 2 2013;3:2651.

58. Northcott PA, Buchhalter I, Morrissy AS, et al. The whole-genome landscape of medulloblastoma subtypes. Nature. Jul 19 2017;547(7663):311–317.

59. Wala JA, Shapira O, Li Y, et al. Selective and mechanistic sources of recurrent rearrangements across the cancer genome. bioRxiv. 2017.

60. Gao J, Aksoy BA, Dogrusoz U, et al. Integrative analysis of complex cancer genomics and clinical profiles using the cBioPortal. Sci Signal. Apr 2 2013;6(269):pl1.

61. Finn RS, Dering J, Conklin D, et al. PD 0332991, a selective cyclin D kinase 4/6 inhibitor, preferentially inhibits proliferation of luminal estrogen receptor-positive human breast cancer cell lines in vitro. Breast Cancer Res. 2009;11(5):R77.

